# Two microbiota subtypes identified in Irritable Bowel Syndrome with distinct responses to the low FODMAP diet

**DOI:** 10.1101/2021.05.14.444142

**Authors:** Kevin Vervier, Stephen Moss, Nitin Kumar, Anne Adoum, Meg Barne, Hilary Browne, Arthur Kaser, Chris Kiely, B Anne Neville, Nina Powell, Tim Raine, Mark D. Stares, Ana Zhu, Juan De La Revilla Negro, Trevor Lawley, Miles Parkes

**Affiliations:** The Wellcome Trust Sanger Institute; Addenbrooke’s Hospital; Wellcome Trust Sanger Institute; Wellcome Sanger Institute; Royal North Shore Hospital

**Keywords:** Microbiome, Irritable Bowel Syndrom, low FODMAP diet

## Abstract

**Objective:** Reducing FODMAPs can be clinically beneficial in IBS but the mechanism is poorly understood. We aimed to detect microbial signatures that might predict response to the low FODMAP diet and assess whether microbiota compositional and functional shifts could provide insights into its mode of action.

**Design:** We used metagenomics to determine high-resolution taxonomic and functional profiles of the stool microbiota from IBS cases and household controls (n=56 pairs) on their usual diet. Clinical response and microbiota changes were studied in 41 pairs after 4 weeks on a low FODMAP diet.

**Results:** Unsupervised analysis of baseline IBS cases pre-diet identified two distinct microbiota profiles, which we refer to as IBS^P^ (pathogenic-like) and IBS^H^ (health-like) subtypes. IBS^P^ microbiomes were enriched in Firmicutes and genes for amino acid and carbohydrate metabolism, but depleted in Bacteroidetes species. IBS^H^ microbiomes were similar to controls. On the low FODMAP diet IBS^H^ and control microbiota were unaffected, but the IBS^P^ signature shifted towards a health-associated microbiome with an increase in Bacteroidetes (p=0.009), a decrease in Firmicutes species (p=0.004) and normalization of primary metabolic genes. The clinical response to the low FODMAP diet was greater in IBS^P^ subjects compared to IBS^H^ (p = 0.02).

**Conclusion:** 50% of IBS cases manifested a ‘pathogenic’ gut microbial signature. This shifted towards the healthy profile on the low FODMAP diet; and IBS^P^ cases showed an enhanced clinical responsiveness to the dietary therapy. The effectiveness of FODMAP exclusion in IBS^P^ may result from the alterations in gut microbiota and metabolites produced. Microbiota signatures could be useful as biomarkers to guide IBS treatment; and investigating IBS^P^ species and metabolic pathways might yield insights regarding IBS pathogenic mechanisms.

Significance of this study

What is already known on this subject?

- IBS subjects often respond to a low FODMAP diet.
- The gut microbiota has been implicated in IBS.
- The microbiota in IBS subjects may change with diet.

What are the new findings

- We were able to stratify patients with IBS according to their gut microbiota species and metabolic gene signatures.
- We identified a distinct gut microbiota subtype with an enhanced clinical response to a low FODMAP diet compared to other IBS subjects.

How might it impact on clinical practice in the foreseeable future?

- The potential development of a microbiota signature as a biomarker to manage IBS cases with a low FODMAP diet recommendation.
- If the bacteria represented in the IBS^P^ subtype are shown to play a pathogenic role in IBS, perhaps through the metabolic activity this provides a target for new therapies and an intermediate phenotype by which to assess them.

## Introduction

Irritable Bowel Syndrome (IBS) affects 10-15% of the population worldwide[1]. It impacts quality of life[2] and incurs significant health economic cost[3]. The pathophysiology of IBS includes changes in visceral nerve sensitivity[4], intestinal permeability[5] and psychological factors[6]. Several lines of evidence suggest the gut microbiome as a key aetiological factor in IBS. For example, there is a six-fold increased risk of developing IBS following an episode of infective gastroenteritis[7], probiotics and dietary intervention can reduce the symptoms [8, 9] and faecal transplantation has reported efficacy in treating IBS[10]. Recent studies using 16S ribosomal RNA profiles (low resolution taxonomic profiling to genus level and no functional inference) have suggested an altered gut microbiota in IBS subjects compared to controls. Although the findings of earlier studies vary significantly recent studies more consistently indicate a reduction in Bacteroidetes in IBS cases vs controls[11-13]. However, mechanisms linking the gut microbiota and IBS symptoms remain poorly understood.

IBS symptoms can be treated with low fibre diets to reduce the colonic microbial fermentation that produces hydrogen and methane, leading to bloating[9]. More recently diets avoiding fermentable oligosaccharides, disaccharides, monosaccharides and polyols (FODMAPs) have demonstrated efficacy [14-17]. The mechanisms are debated[18], but potentially involve modulation of microbiota composition and metabolite production[19].

The low FODMAP diet is challenging for many patients to follow, reduces various ‘healthy’ foodstuffs (pulses, particular fruits and vegetables), and its long-term consequences on health are unknown. Thus there is a recognised need to better understand how low FODMAP diets work[20], and ideally identify biomarkers that predict response.

In order to accurately link changes in gut microbiota structure with diet, including low FODMAP diets, detailed taxonomic profiling and quantification of microbial abundance is required. The gut microbiota of healthy adults is diverse, dominated by hundreds of bacterial species from the Bacteroidetes and Firmicutes phyla, with fewer species from Actinobacteria and Proteobacteria [21]. It is shaped by diet and impacts immunity, metabolism and cognition[22, 23]. While 16S rRNA studies have provided valuable insights into the gut microbiota and IBS, they cannot achieve taxonomic resolution to species level. Techniques of microbial culture and metagenomic sequencing now enable detailed taxonomic and functional characterisation [24].

The aim of the present study was to identify a biomarker of response to the low FODMAP diet and gain insights into microbial changes underlying treatment success using high resolution metagenomic and functional analysis of subjects with IBS and household controls before and while on a low FODMAP diet.

## Materials and methods

### Subjects

A prospective single centre case control study recruited participants from 2016 to 2019. We included adults (18-68 years of age) meeting the Rome IV criteria[25] for diarrhoea-predominant or mixed type IBS (IBS-D and -M respectively) with respective household controls. Subjects were recruited from outpatient clinics at Cambridge University Hospital in the UK and via a social media campaign.

We excluded cases with other gastrointestinal diseases, pregnancy, those already following a restrictive diet, and those taking probiotics or other medications within one month that could potentially modify the gut microbiota such as antibiotics, proton pump inhibitors, colonoscopy bowel preparation or metformin[26].

#### Ethics

approval was provided by Cambridge Central Research Ethics Committee reference 15/LO/2128.

Study procedures are summarised in Figure 1. Participants were assessed at baseline by a consultant. Three subsequent study visits were supervised by a dietitian when symptom severity scores were captured using the IBS Severity Scoring System (IBS-SSS)[27] and 7 day food intake diary used to assess FODMAP intake.

**Figure 1.**
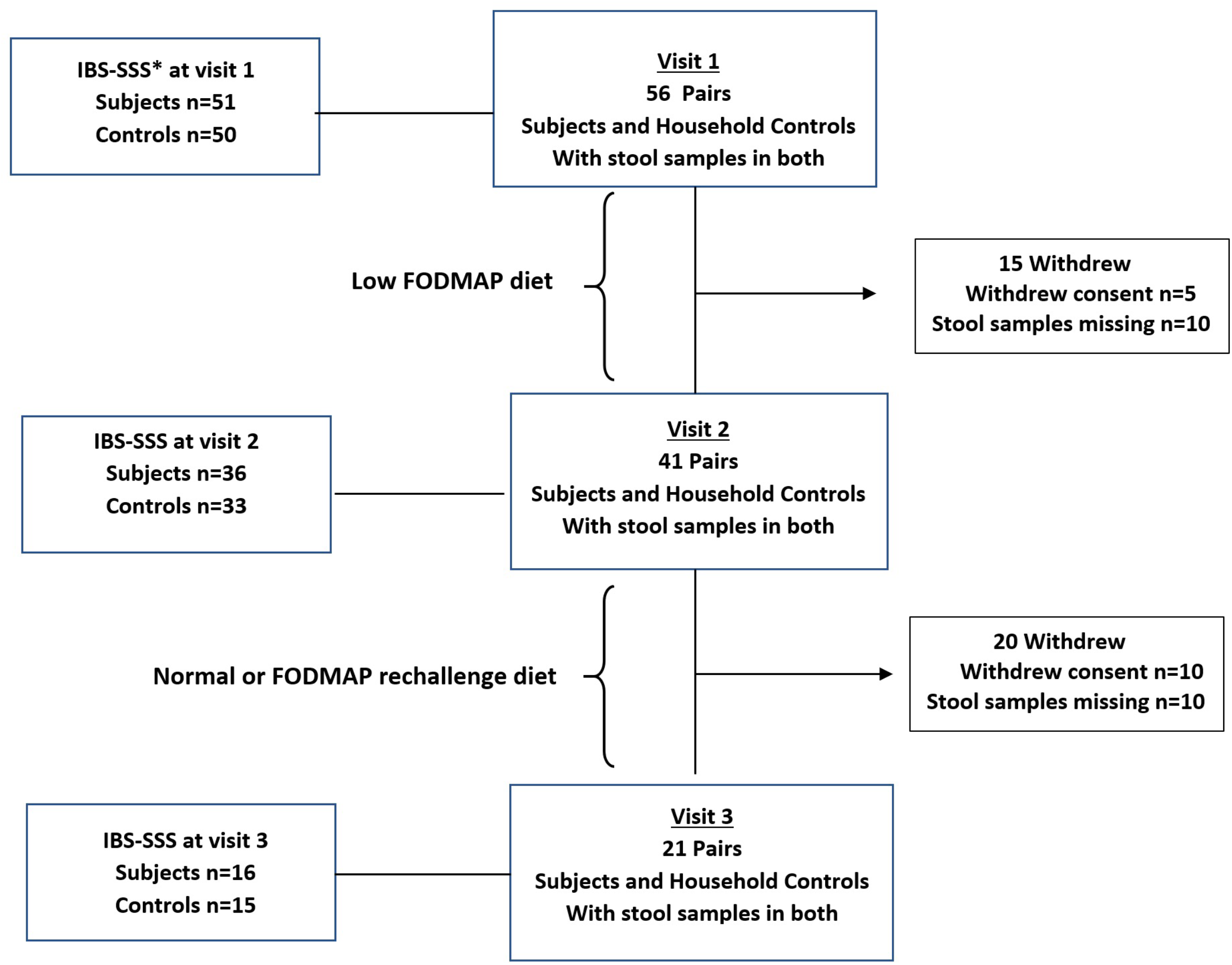
Flowchart for IBS microbiome study: number of pairs of IBS subjects and each of their household controls providing stool samples at visits 1 – 3. IBS-SSS* – Irritable Bowel Syndrome Severity Scoring System

### Stool samples

Participants and their household controls were asked to provide a stool sample at visit 1 while on their usual diet, after 4 weeks on a low FODMAP diet (at visit 2) and 12 weeks following FODMAP rechallenge in IBS subjects improving on the diet (visit 3, to identify individual FODMAP triggers), or a return to usual diet in IBS subjects not improving with the diet and in all household controls. Samples were sealed and immediately placed in the participant’s home freezer then courier transferred on dry ice to the Wellcome Sanger Institute within 48 hours for storage at -80°C prior to processing. DNA was extracted using the MP Biomedicals™ FastDNA™ SPIN Kit for Soil.

### Metagenomic Sequencing

To profile the taxonomic composition of the stool samples from cases and controls we performed shotgun metagenomic sequencing using the Illumina Hi-Seq 4000 platform (read length 150bp, 450bp fragment size, average 12 million paired-end reads). Raw sequencing data were deposited under ENA Study Accession Number: XXXXXX. Paired-end read files were classified using a Kraken2 bespoke database containing 3,000 high-quality assemblies from 784 species associated with the human gut microbiome. Bracken[28] was applied to obtain refined species-level metagenomic profiles. An average of 10.2 million sequencing reads were classified at species rank by our platform, corresponding to a read classification rate of 86% (sup Fig 1). No difference in assigned read counts was observed between cases and controls (Wilcoxon P=0.7). Statistical analysis was performed using R language [29]. Taxonomic profiles were normalized using center log ratio (CLR) transform[30] after estimating zero values using the *cmultRepl* function from *zCompositions* R package [31].

For an unsupervised analysis to identify sub-populations of IBS cases, we applied a k-means clustering algorithm to CLR-transformed taxonomic profiles from baseline IBS case and household control samples. The optimal k value was obtained by minimizing within-group sum of squared metrics using the *decostand* function from the *vegan* R package[32].

A comparison of alpha diversity between groups was performed using a paired Wilcoxon test.

We used Aitchison distance[30] which is the Euclidean distance of the CLR transformed profiles to estimate beta diversity between samples. The significance of beta diversity difference was estimated using PERMANOVA test[33].

Associations between cluster assignment and clinical metadata were sought using Fisher’s exact test on the contingency table or Mann-Whitney test when appropriate (Table 1). We tested for correlation between microbial abundance and metadata (including IBS cluster and timepoint) by applying generalized linear mixed models (GLMMs) using MaAsLin2 software[34]. Random effects were modelled by matching IBS subject and household control. Vignette and source code for the analysis are available at http://github.com/kevinVervier/IBS.

**Table 1.** Demographic characteristics of the 56 IBS subjects according to the cluster separation based on the differences in the microbiome. Fisher’s test was applied on categorical variables (Sex, IBSD, IBSM, post infectious IBS, medications, smoking status), Wilcoxon test was applied on continuous variables (age, BMI, IBS-SSS, alcohol intake) to estimate statistical significance of the difference between groups. IBS-SSS* – Irritable Bowel Syndrome Severity Scoring System

|  | Cluster 1 (n=28) | Cluster 2 (n=28) | P-value |
| --- | --- | --- | --- |
| Female(%) | 22 (79) | 19 (68) | 0.84 |
| Age (mean ± SD) | 37.4 ± 12.5 | 39.9 ± 14.4 | 0.54 |
| BMI (mean ± SD) | 29.4 ± 7.7 | 26.5 ± 5.4 | 0.08 |
| IBSD (%) | 12 (43) | 11 (39) | 1 |
| IBSM (%) | 16 (57) | 17 (61) | 1 |
| Post Infectious IBS (%) | 8 (28) | 6 (21) | 0.77 |
| Median IBS-SSS | 302 (n=24) | 249 (n=27) | 0.17 |
| Antidepressants (%) | 6 (21) | 3 (11) | 0.48 |
| PPIs / H2RAs (%) | 1 (4) | 0 (0) | 1 |
| Smokers (%) | 4 (14) | 1 (4) | 0.36 |
| Median Alcohol Intake (U/Week) | 3.5 (n=24) | 0.5 (n=26) | 0.13 |

2,754 high quality human gastrointestinal genomes (Attached Table 1) were downloaded from Human Gastrointestinal Bacteria Culture Collection[24], Culturable Genome Reference[35] and National Center for Biotechnology Information (NCBI). Maximum-likelihood trees were generated using FastTree v2.1.10[36] with default parameters, and protein alignments were produced by GTDB-Tk v1.3.0[37] with the *classify_wf* function and default parameters. Trees were visualized and annotated with Interactive Tree Of Life (iTOL) v5[38].

### Functional metagenomic and genomic analysis

Functional profiling on each metagenome was conducted using HUMAnN3[39] with default parameters to quantify MetaCyc pathways[40]. Pathway enrichment was performed using MaAsLin2[34] (threshold at *q*-value < 0.1). Enriched pathways were classified in broad categories using the MetaCyc database.

To identify the genes present in an enriched MetaCyc pathway in a reference genome, we first collected the protein sequence corresponding to each gene in each pathway from the Metacyc database and UniProt[41]. BlastP[42] was then performed for each of these protein sequences against a protein database based on 544 genomes with a cut-off E-value of 1e-10. This genome collection of 544 genomes includes 420 genomes (56 species) of IBS associated bacteria representing cluster IBS^P^ and 124 genomes (34 species) of health-associated bacteria representing cluster IBS^H^ (see below for IBS^P^ / IBS^H^ description). Gene enrichment was calculated using one-sided Fisher’s exact test with p-value adjusted by Hochberg method.

## Results

### Cohort Summary

The cohort is summarised in Figure 1. Among cases, there was female predominance (73%) and IBS-M was the commonest subtype (59%). Fourteen cases (25%) reported symptom onset after an episode of gastroenteritis. The median IBS-SSS at baseline in the 56 cases was 272, with 45 cases (88.2%) scoring moderate (IBS-SSS>175 – n= 25) or severe (IBS-SSS> 300 – n=20). In controls the median IBS-SSS score was 7.5 (range: 0-196). Mean age of subjects was 38.7 (range 18-68) and controls 44.6 (range 18-74).

### Comparison of gut microbiota from IBS cases and household controls

Metagenomic sequencing was carried out on 234 stool samples followed by reference genome mapping of sequence reads [24]. Our inclusion of household controls reduced confounding by environmental exposures (pets, prevailing diet, hygiene regime) and is important as gut microbes can frequently transmit between co-habiting humans[43]. Indeed, we observed that samples coming from the same household had a more conserved microbiota composition compared to the overall variability between all cases and all controls (sup Fig 2C, Wilcoxon p = 6.02E-05). To account for this potential confounder in subsequent analyses we applied pairwise comparisons where possible.

We first focused on understanding the compositional variation in bacterial species to identify potential pathogenic imbalances in IBS case gut microbiomes. We measured alpha diversity in baseline samples using the Chao1 index for the number of species (richness) and the Shannon index for the relative abundance of different species (evenness). The richness was not lower in IBS cases (Wilcoxon p=0.12) (sup Fig 2A), but the lower evenness suggested a more imbalanced species abundance distribution in favour of fewer bacterial species in IBS cases compared to controls (p=0.0092; sup Fig 2B). We also measured beta diversity between baseline microbiota samples using Aitchison distance and observed significantly more taxonomic variability within IBS cases compared to controls (sup Fig 2C, Wilcoxon p=1.3E-79).

### Stratification of IBS patients based on gut microbiota compositional subtypes

The high variability in diversity observed within baseline microbiomes from IBS cases warranted exploration of possible stratification by microbiome profile, to identify distinguishing signals that went undetected during our initial analysis. We therefore performed unsupervised data clustering designed to identify microbiota subtypes in baseline samples from the 56 pairs of cases and household controls. Importantly, this analysis revealed two distinct microbiota taxonomic clusters (Fig.2A) with 28 cases assigned to each. Such clustering was not seen in household controls. Microbiome compositions in cluster 2 cases were similar to healthy controls whereas the cluster 1 cases were clearly separated. Compared to the overall variability previously observed across all IBS cases, microbiota diversity within each cluster was more conserved (sup Fig 3, Wilcoxon p=2.2E-08). We found no significant difference in age, gender, BMI, subtype of IBS, post-infectious IBS or concomitant medications between the two clusters (Table 1). Baseline symptom severity scores appeared modestly higher in cluster 1 than cluster 2 (median IBS-SSS=302 vs 249), but this was not statistically significant (Wilcoxon p=0.17).

**Figure 2.**
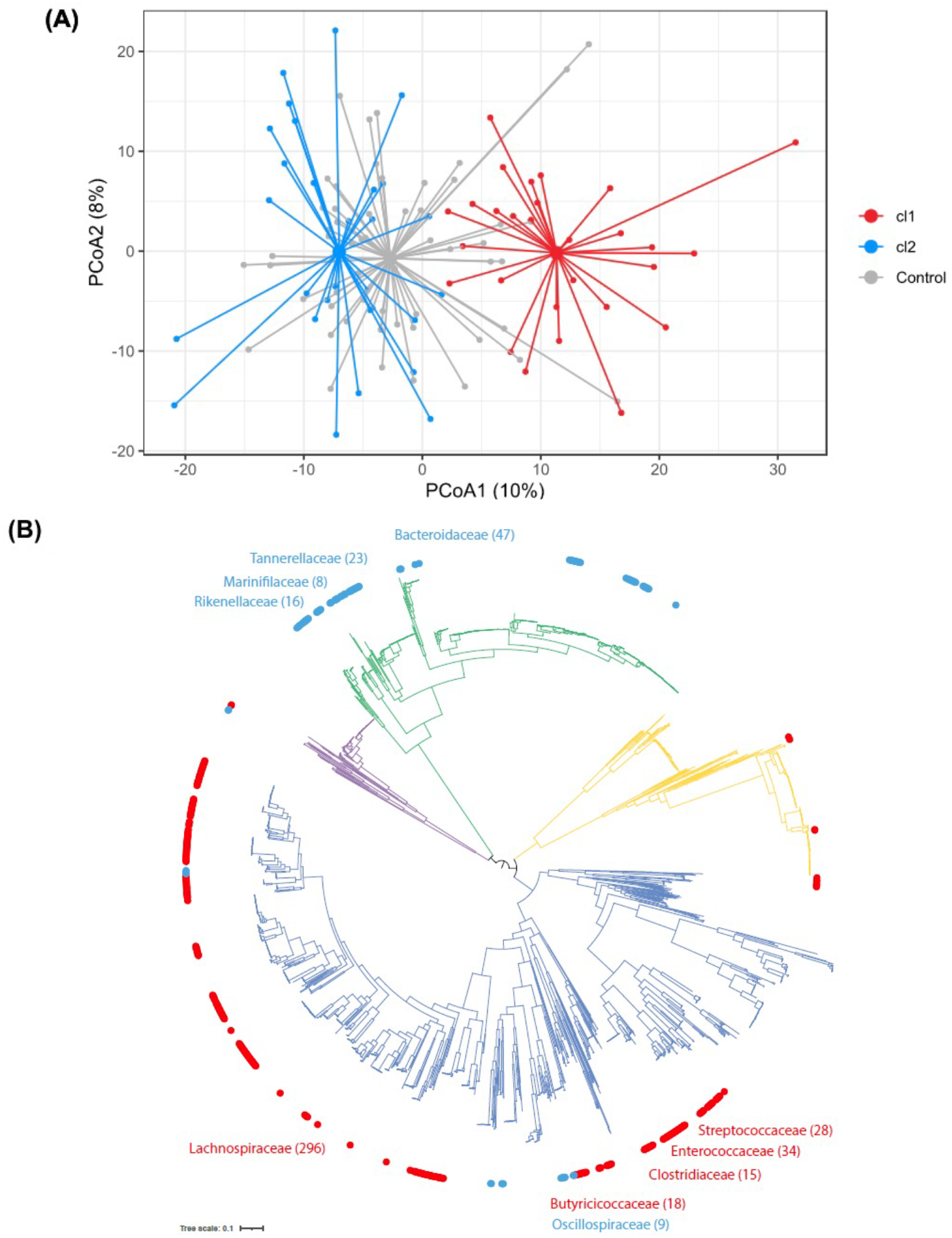
analysis of diversity of microbiota profiles (A) Beta diversity analysis of IBS cases and healthy controls: principal coordinate analysis for the two first components identifies two distinct clusters among cases, described as cluster 1 (cl1, red) and cluster 2 (cl2, blue). Overall dispersion of healthy participants is represented in grey. Variance explained by PC1: 14%, PC2: 9%. (B). Phylogenetic tree of 2754 human gut bacterial isolates generated using the 120 core genes. Outer circle distinguishes bacteria abundant in cl1 (red; n = 420 genomes) and bacteria abundant in cl2 (blue; n = 124 genomes) shown. Top5 prevalent families in each cluster are named. Branch colour distinguishes bacterial phyla belonging to Actinobacteria (yellow; n⍰= ⍰363 genomes), Bacteroidetes (green; n⍰= ⍰675 genomes), Firmicutes (darkblue; n⍰=⍰1562 genomes), and Proteobacteria (purple; n⍰=⍰154 genomes) shown.

The number of bacterial species (richness) appeared modestly lower in cluster 1 cases compared to cluster 2 microbiomes (Wilcoxon p=0.033), but no such difference was observed between respective controls (Wilcoxon p=0.57) (sup Fig 4A). Cases and controls from the same cluster show comparable richness (cluster 1 p=0.073, cluster 2 p=0.69). Shannon diversity (evenness) was clearly lower in IBS cluster 1 compared to cluster 2 cases (p=0.0002), but this difference was not seen between respective controls (p=0.078) (sup Fig 4B). Cases from cluster 1 had a lower evenness when compared to their household controls (p=0.0029), while this was not observed for cluster 2 (p=0.41). Overall, our findings suggest that cluster 1 case microbiomes are depleted in bacterial species and skewed towards specific bacteria compared to cluster 2.

Read abundance analysis identified distinct differences between bacterial species in the two IBS subtypes at baseline (MaAsLin2 q-value < 0.1; Attached Table 2). A total of 87 species were identified as significantly differentially abundant between the two IBS subtypes (56 up in cluster 1 and 31 up in cluster 2), and not observed between corresponding household controls. In IBS cluster 1 we observed a significant increase of bacteria from the Firmicutes phylum including known human pathogens (*Clostridium difficile, Paeniclostridium sordellii, Clostridium Perfringens, Streptococcus anginosis*) (sup Fig 4C) and a significant depletion of multiple *Bacteroides* and *Parabacteroides* species (sup Fig 4D). Phylogenetic analysis showed a clear distinction between the dominant species from the Firmicutes phylum in cluster 1 and the dominant species from the Bacteriodetes phylum in cluster 2 (Fig.2B). However, we did not observe a significant difference in abundance for these two phyla between groups (MaAsLin2 q-value: Firmicutes: 0.2, Bacteroidetes: 0.78) suggesting differences in a subset of species rather than an overall Firmicutes/Bacteroidetes imbalance.

Thus, we identified IBS subtypes with distinct microbiota signatures and clinical features at baseline: cluster 1 contained lower bacterial diversity, was depleted in commensal species from the Bacteroidetes phylum and enriched in species from the Firmicutes phylum, including human pathogens; and cluster 2 was indistinguishable from healthy household controls. We refer to cluster 1 as IBS^P^ microbiome type for its pathogenic properties and cluster 2 as IBS^H^ microbiome type due to its similarity to healthy household controls.

### Enrichment of primary metabolism genes in gut microbiomes of IBS^P^ patients

Bacterial species from the Bacteroidetes and Firmicutes phyla are evolutionarily and physiologically distinct, and contribute different core functions to the gut microbiome. Therefore, we reasoned that the functional capacity of IBS^P^ microbiomes may contribute to IBS symptoms. To identify functional differences between the microbiomes of the two IBS subtypes, we performed an analysis of the functional capacity encoded in the metagenomes of baseline samples of IBS^P^ and IBS^H^ patients. This analysis was independent of the previous taxonomic analysis. We found a significant enrichment of 109 functional pathways and significant depletion of 13 functional pathways in IBS^P^ microbiomes compared to IBS^H^ microbiomes (Attached Table 3). Further functional classification indicated that the majority of enriched pathways in IBS^P^ microbiomes (78.7%) could be classified to 5 major functional categories related to primary metabolism (Figure 3).

**Figure 3.**
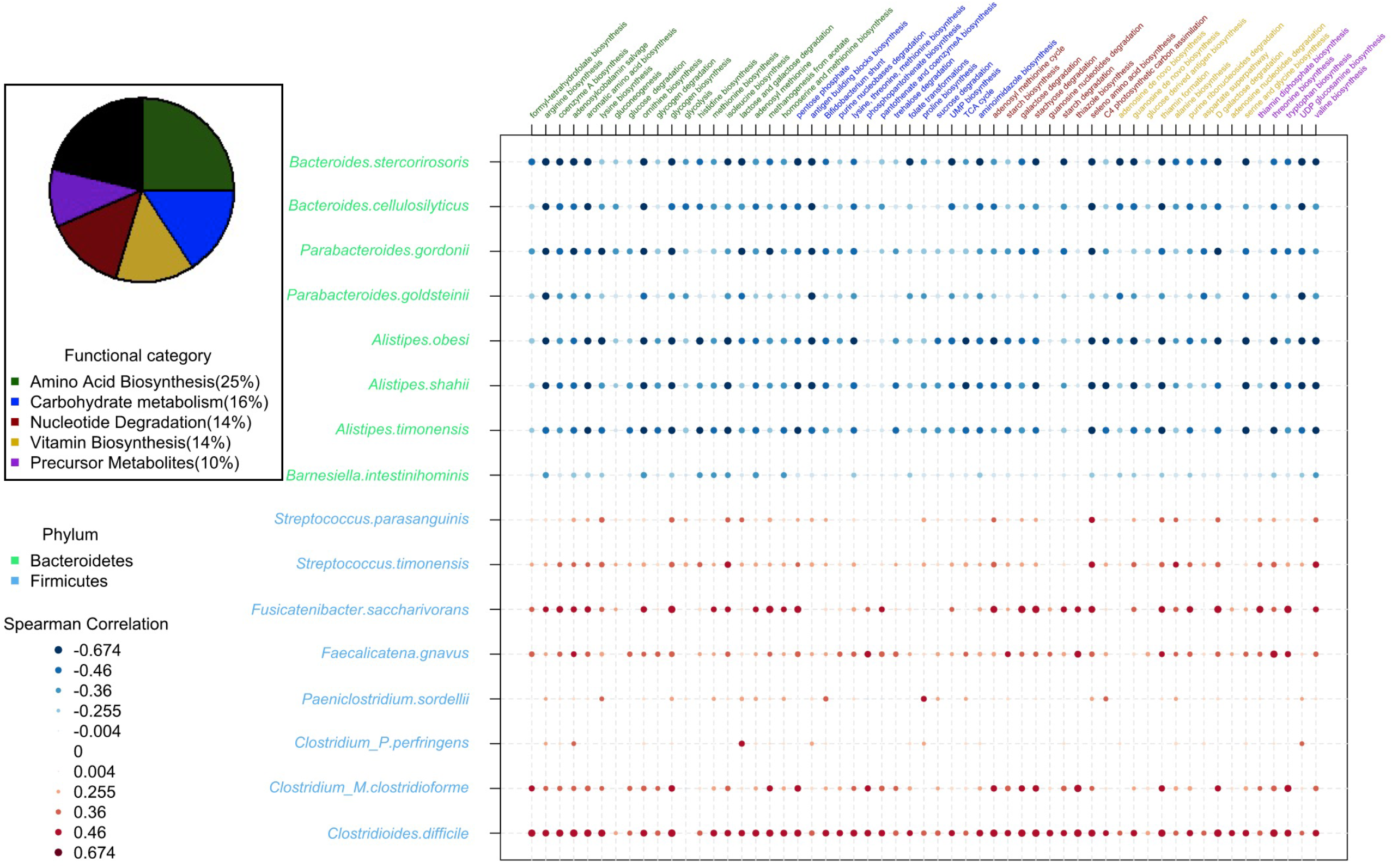
Taxonomic and functional characterization of IBS^P^ subjects baseline microbiome. Pie chart indicates the distribution of pathways identified as significantly enriched in IBS^P^ subjects at baseline and coloured according to their MetaCyc functional category. A selection of candidate pathways are represented in columns (colored as in the piechart). Species significantly different in abundance between IBS^P^ and IBS^H^ subjects are represented in rows and coloured by phylum (Bacteroidetes in green and Firmicutes in blue). For each combination of pathway and species, Spearman correlation on their respective abundance is reported (from strongly positive in red to strongly negative in blue).

Since amino acid biosynthesis (25%) and carbohydrate metabolism (15.7%) were the two major functional categories that separate IBS^P^ and IBS^H^ cases, we next performed a targeted functional enrichment analysis in IBS^P^ microbiomes at the species level. For amino acid biosynthesis this identified significant enrichment of genes involved in biosynthesis of tryptophan, threonine and histidine (sup Figure 5). Equivalent analysis of carbohydrate metabolism identified significant enrichment of genes involved in lactose metabolism, fructose metabolism, and trehalose metabolism, and biosynthesis of two short chain fatty acids (SCFA): butyrate and propionate (sup Figure 6).

Our results suggest specific functions involved in amino acid biosynthesis and metabolism of simple dietary sugars are distinct features in bacteria of the IBS^P^ cluster at baseline, which are underrepresented in bacteria of the IBS^H^ cluster. Correlating the compositional (Figure 2B) and functional (Figure 3) features identified a subset of candidate species associated with the IBS^P^ cluster (Figure 3) and enriched in significant pathways. A strong positive correlation was observed between the abundance of these pathways and abundance of the bacterial species with known pathogenic capabilities (*C. difficile, P. sordellii, C. perfringens*) and a pathobiont associated with ulcerative colitis (*Faecalicatena gnavus*, previously named *Ruminococcus gnavus[44]*). Commensal species depleted in IBS^P^ patients did not encode these pathways.

### Low FODMAP dietary intervention corrects IBS^P^ microbiomes

A total of 41 IBS cases and their household controls followed a low FODMAP diet for 4 weeks and provided a stool sample while on the diet. There was a significant reduction in the IBS-SSS on and after the completion of the low FODMAP diet (mean IBS-SSS pre-diet = 278, on diet = 128, post diet = 117) (Figure 4A). A decrease in IBS-SSS scores was seen on the low FODMAP diet in patients harbouring IBS^P^ and IBS^H^-type microbiomes (Figure 4B) but was more pronounced in IBS^P^ patients (Δ IBS-SSS in IBS^P^ =194 vs IBS^H^ =114; p=0.02) (Figure 4C).

**Figure 4.**
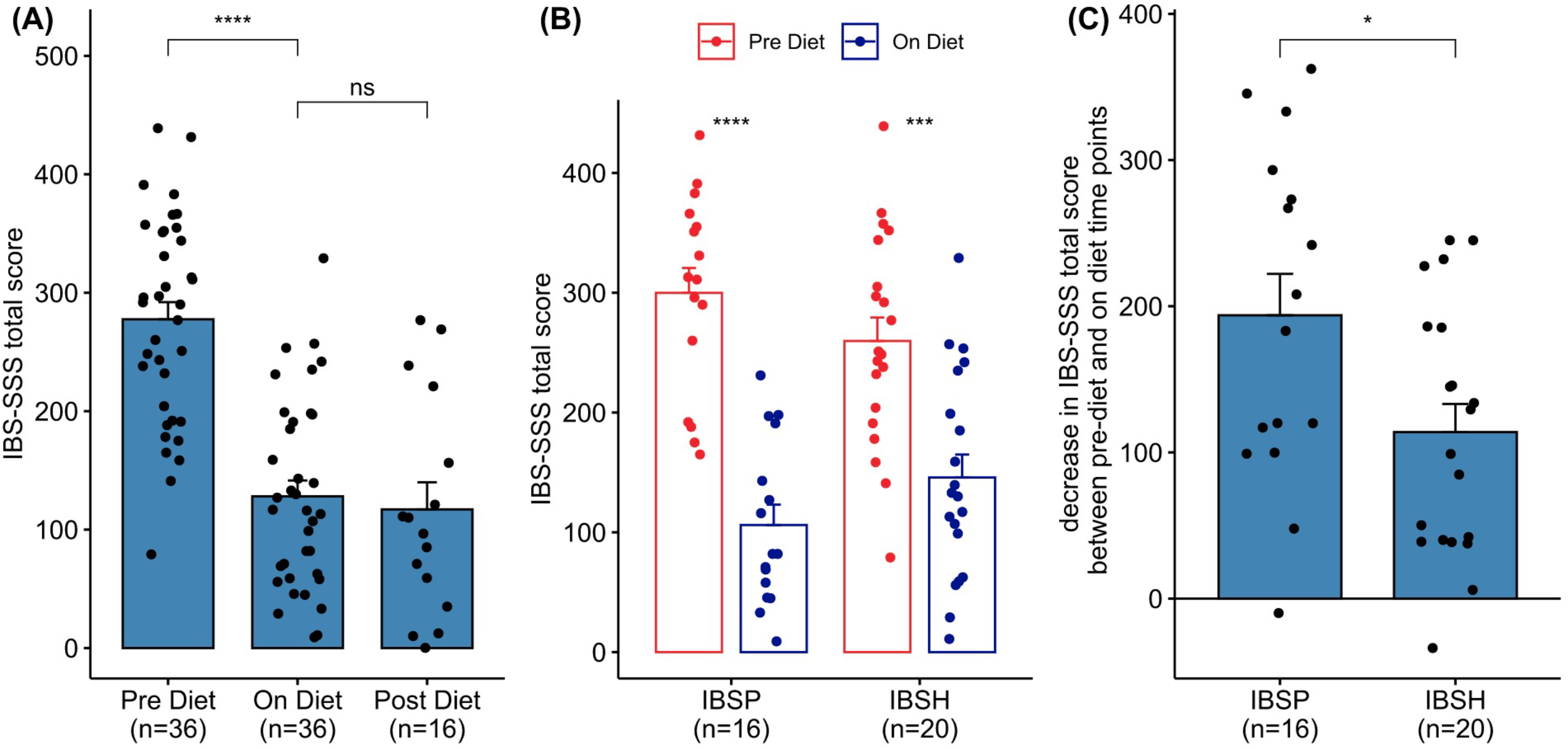
Clinical response in 36 subjects undergoing dietary intervention and providing IBS-SSS. (A) response for combined IBSP and IBSH subjects, also includes IBS-SSS in 16 subjects at visit 3. (B) response pre and on diet according to the microbiota cluster pre diet. (C) Change in IBS-SSS from pre-diet value to on diet value for patients in each cluster. Paired Wilcoxon test was used to estimate statistical significance of the difference between groups (****: p<0.0001, ***: p<0.001, *:p<0.05, ns: p>0.05). Bar height shows mean value, error bars show standard error. IBS-SSS* – Irritable Bowel Syndrome Severity Scoring System

Comparison of taxonomic profiles between baseline (pre-diet) stool samples and those obtained while on the low FODMAP diet for four weeks revealed a significant shift in the microbiota composition of IBS^P^ cases but not IBS^H^ cases nor healthy controls (Figure 5A). Compared to the differences seen between IBS^P^ and IBS^H^ at baseline, beta diversity analysis showed the microbiome profiles from IBS^P^ cases became more similar to those seen in IBS^H^ cases and healthy controls while on the low FODMAP diet. This was apparent as a decreased variability in microbiome composition within all IBS cases (IBS^P^ + IBS^H^ combined) on diet compared to pre-diet (Supp. Fig. 7, Wilcoxon test p=1E-19). Within both IBS^P^ and IBS^H^ cases it was also evident that the diet produced a greater shift in microbiota composition in IBS^P^ compared to IBS^H^, with a bigger distance between sample profiles from the same case at the two timepoints (baseline and on-diet) (Supp. Fig. 7, Wilcoxon test p=0.03).

**Figure 5.**
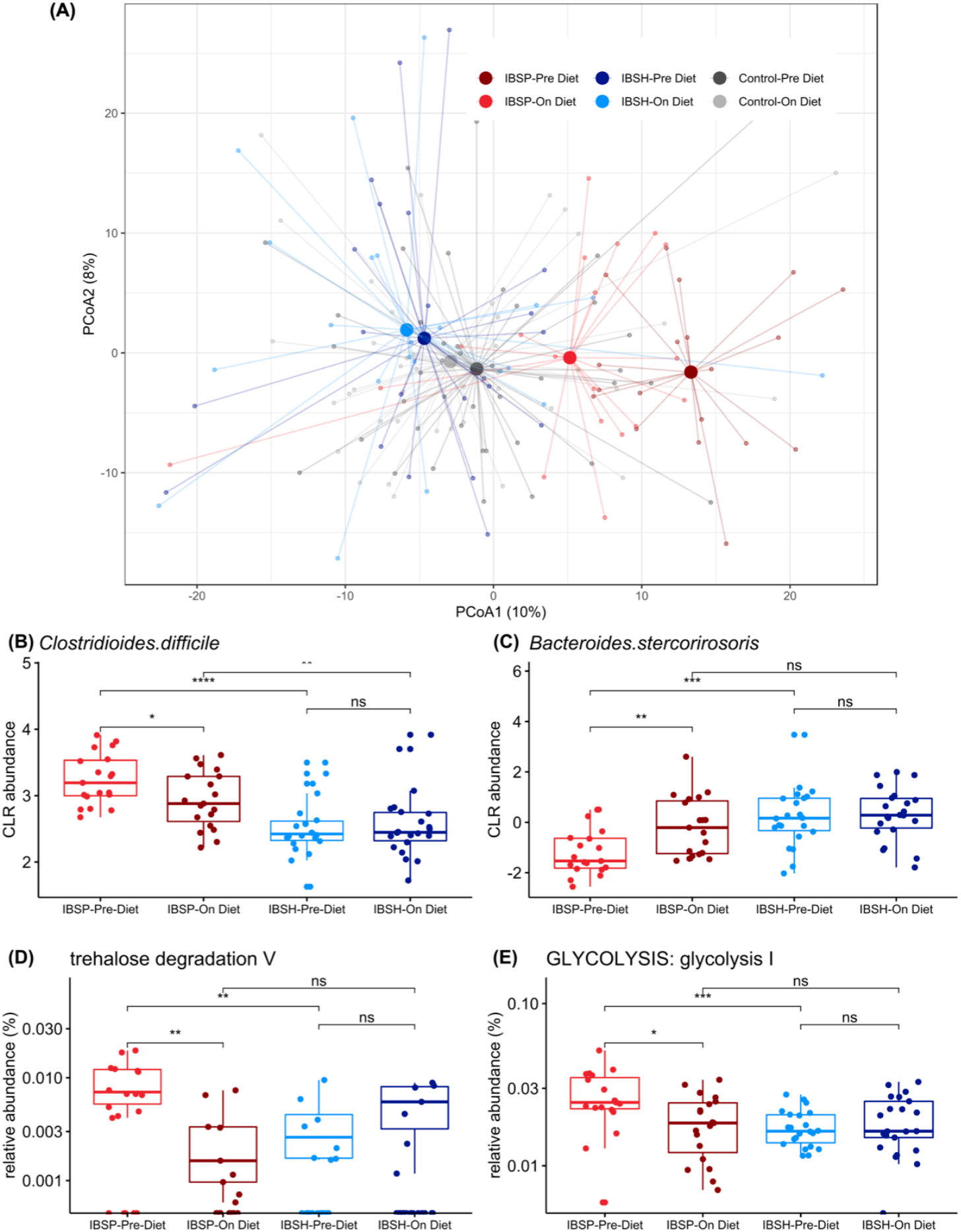
Microbiome beta-diversity before and during diet intervention. (A) Principal coordinate analysis of IBS cases separated into two clusters showed a diet-triggered shift in IBS^P^ (red) only with respect to IBS^H^ subjects (blue) and healthy controls (grey). (B,C) Impact of diet intervention on taxonomic abundance. Linear mixed models identified differentially abundant species between IBS^P^ and IBS^H^ cases pre-diet and on diet. Center logratio (CLR) transformed abundances for representative species are shown. (B) Pathobiont species, such as C. difficile, become less abundant in IBS^P^ during diet intervention, (C) members of Bacteroides genus become more abundant in IBS^P^ during diet intervention. (D,E) Impact of diet intervention on pathway abundance. Relative abundances for representative pathways are shown. (D) degradation of the FODMAP trehalose became less abundant in IBS^P^ during diet intervention, (E) glycolysis became less abundant in IBS^P^ during diet intervention. Wilcoxon test was used to estimate statistical significance of the difference between groups (****: p<0.0001, ***: p<0.001, **: p<0.01, *:p<0.05, ns: p>0.05). Box and whiskers show, median and interquartile range.

Diet intervention shifted the taxonomic composition of IBS^P^ cases by increasing Bacteroides levels (*B. cutis, B. stercorirosoris*), and decreasing pathobiont levels (including *C. difficile, Streptococcus parasanguinis, Paeniclostridium sordelli*) towards those seen in IBS^H^ (Fig. 5 B-C) and household controls (Supplementary Figure 8). The functional profile of the IBS^P^ microbiome was also impacted by the diet intervention, for example producing a decrease in degradation of the FODMAP trehalose (Fig. 5D) and a decrease in glycolysis to levels comparable to those in IBS^H^ patients and healthy controls (Fig. 5E).

After the low FODMAP diet ended participants returned to a normal diet, albeit with cases excluding foods identified as triggering their symptoms. After 3 months there was no significant shift in the microbiota diversity of the cases in the two clusters compared to while on full dietary restriction (Supp. Fig 9, PERMANOVA p=0.998) and no significant change in the abundance of any bacterial taxa between these timepoints. Thus, the shift in the IBS^P^ microbiota to a heathy profile appeared stable for at least 3 months and correlated with continuing symptomatic well-being (Fig. 4A).

## Discussion

We defined two gut microbiome subtypes in IBS cases with distinct signatures based on species and encoded microbial functions, and differential clinical responses to a low FODMAP diet intervention. Although the early IBS microbiome literature is rather inconsistent regarding taxa implicated and the presence of subtypes [11], our work is congruent with the observations of Jeffery *et al*.[12, 13] who used shotgun, 16S rRNA gene microbiome profiling, and metabolomics to provide evidence of IBS microbiome subtypes identifying Lachnospiraceae species and enrichment in amino acid biosynthesis. Not only do our results replicate this stratification within IBS in a larger cohort, but being based on shotgun metagenomics data they benefit from both greater taxonomic resolution - identifying an increase in selected Firmicutes species and depletion of Bacteroidetes species in one subgroup - and the ability to analyse the functions encoded in the microbiome. Furthermore, the dietary intervention allowed us to characterize the clinical responses of each patient subtype; and inclusion of household controls, following the same dietary intervention, was a unique feature of our study designed to correct known confounding environmental effects[45].

We refer to the IBS microbiome subtypes as IBS^p^ (pathogenic) and IBS^H^ (healthy). Overall, 75% of IBS cases in our study improved on a low FODMAP diet as measured by a decrease in IBS-SSS but higher levels of symptom response were seen in cases with IBS^P^ compared to IBS^H^ microbiomes. IBS^P^ microbiomes were notably different from the microbiome of IBS^H^ cases and healthy household controls, with an enrichment of distinct bacterial species and gene families seen in IBS^P^ that allows us to propose potential pathogenic mechanisms.

Within the dysbiotic IBS^P^ microbiomes we saw a significant enrichment of a broad range of evolutionarily distinct Firmicutes species, including known human pathogens (*Clostridium difficile, C. sordellii and C. perfringens*), a pathobiont associated with ulcerative colitis (*Faecalicatena gnavus*, previously named *Ruminococcus gnavus*[44]) and known gut species not previously identified as human pathobionts (*C. clostridioforme and Fusicatenibacter saccharivorans*). Interestingly, we also saw an enrichment in IBS^P^ microbiomes of the lactic acid bacteria *Streptococcus parasanguinsis* and *S. timonensis*, that are usually found in the oral cavity.

IBS^P^ microbiomes are enriched in genes and pathways involved in metabolising simple sugars that are recognised FODMAPs commonly found in dairy products (lactose), fruit (fructose) and food additives (trehalose, lactose and fructose). Lactose and fructose are known triggers of IBS so our analysis provides a list of candidate bacteria for further investigation (sup Figure 5). Interestingly, trehalose is found in mushrooms (excluded in the low FODMAP diet) and was introduced as a food additive in the 1990s, since when specific lineages of *C. difficile* have evolved to avidly metabolize it and in so doing increase their abundance [46]. Trehalose could trigger IBS symptoms by fuelling the growth of specific ‘pathogenic’ bacterial species.

Microbial metabolism of hexoses derived from FODMAP carbohydrates produce pyruvate by anaerobic glycolysis in the gut. Pyruvate is a key metabolite that feeds in to short chain fatty acid (SCFA) production[47]. Our pathway analysis (sup Figure 5) predicts that several bacterial species enriched in IBS^P^ microbiomes contain genes for converting pyruvate to butyrate (classical pathway) and/or propionate (acrylate pathway)[48]. Butyrate and propionate are major metabolites in the colon that bind to GPR receptors 41, 43 (propionate) and 109A (butyrate): these short chain fatty acids regulate tryptophan hydrolase gene transcription in enterochromaffin cells facilitating the production of 5 hydroxytryptamine (5HT) from tryptophan; 5HT is postulated as a key agent in the production of IBS symptoms[49, 50]. Moreover, in IBS^P^ microbiomes, we observed an enrichment of genes for tryptophan biosynthesis which would facilitate this mechanism.

We also found enrichment in IBS^P^ microbiomes for the genes coding for the biosynthesis of amino acids including histidine, arginine, ornithine, tryptophan, alanine and threonine (supp Figure 6). Interestingly, Lee et al.[51] found elevated levels of threonine, tryptophan, and phenylalanine, as well as amino acid metabolites cadaverine and putrescine, in stool samples of IBS patients, providing direct evidence of altered amino acid metabolism. Histidine is a precursor to histamine, implicated in the generation of IBS symptoms following its release from mast cells; histamine can itself also activate these cells [42].

Although we detect higher levels of specific pathogens in IBS^P^ microbiomes we have no evidence to suggest they are causing IBS symptoms through known toxin virulence factors. Instead, the data suggest an enrichment of primary metabolic pathways in diverse Firmicutes species. Our analysis indicates a potential for increased production of amino acids; and SCFA through metabolizing FODMAP carbohydrates. It is possible that such metabolites and their derivatives could be noxious at high levels within the colon, or be pathological if produced within the wrong intestinal niche, a type of metabolic virulence, leading to IBS symptoms. One key finding from our work is that IBS^P^ and IBS^H^ microbiomes have distinct bacterial community responses to low FODMAP dietary intervention, providing a basis to define a mode of action. Thus it is possible that removal of the eliciting dietary component starves the pathobionts leading to reduction in their growth and metabolism and a consequent decrease in symptoms, accompanied by an expansion of commensal or symbiotic species leading to a health associated microbiome. Although the number of case/control pairs (n=21) who provided follow-up samples at 12 weeks after rechallenge with FODMAPs was relatively modest, and some continued to exclude specific FODMAP-containing foods, it was interesting to note that both their symptoms (Fig 4A) and microbiomes (sup Fig 9) remained notably stable. This corroborates and perhaps helps to explain the durable benefit that can be seen from a low FODMAP diet.

We observed a differential response of IBS^P^ and IBS^H^ microbiome subtypes to the low FODMAP diet, suggesting that some gut microbiomes are more influenced by dietary interventions. Based on our analysis it is not obvious how or whether IBS^H^ microbiomes contribute to IBS symptoms since they are indistinguishable from household control microbiomes and did not significantly alter in response to the low FODMAP diet. That symptoms in IBS^H^ cases still improved somewhat on FODMAP exclusion suggests either that the response is linked to a non-bacterial component of the microbiome, such as viruses, or is unconnected mechanistically to the microbiota, perhaps instead reflecting a direct effect of dietary constituents and their metabolites on gut neuronal function or osmotic load.

The presence of microbially-defined IBS subtypes with differing responses to dietary intervention has been suggested by some previous studies. In one the microbiome in children responding to a low FODMAP diet appeared enriched at baseline with taxa such as Bacteroides, Ruminococcaceae and Faecalibacterium prausnitzii[14]. In other studies, stool microbial profiles assessed by a commercial kit correlated with differing responses to a low FODMAP diet[52]; and the profile of faecal volatile organic compounds, postulated as reflecting microbiome differences, predicted response to a low FODMAP diet or probiotics[53].

Our study has limitations. The sample size was relatively modest: the strict inclusion criteria, the restriction of concomitant medications and the required participation of household controls needing to follow the low FODMAP diet hindered recruitment. Dietary information was limited to the last week of the interventional phase of the low FODMAP diet: participants could have been tempted to follow a more rigorous diet on the week they had to report their dietary intake. With the design of the study, it was impossible to exclude other factors, apart from diet, that could have impacted the benefit observed, including the psychological impact of being assessed within a research study, the placebo effect that has been described in other studies, and referral bias. Our findings of distinct IBS clusters based on microbiome profiles, the shift on the low FODMAP diet and the clinical responses, should be validated in other populations from different geographical distributions and exposed to different dietary habits.

The identification of a microbial signature ‘biomarker’ that correlates with improved response to a low FODMAP diet may, if validated, allow better stratification and selection of patients likely to benefit from the diet. It also opens the door to trying other therapeutic strategies that manipulate the microbiota in the same direction and achieve the same symptomatic improvement but without the need to undergo the same stringent dietary restrictions. Further, closer study of the implicated microbes may give the opportunity to better understand the interaction between diet, microbiota, metabolites and the human gut-brain axis that leads to the development of IBS symptoms in more than 10% of the world’s population.

## Acknowledgements

We would like to thank the patients and household controls who participated in the study. Also Tracy Papworth, Max Delvincourt, Chris Cederwall and You Yi Hong who contributed to patient identification and recruitment. This research was co-funded by Addenbrooke’s Charitable Trust (ACT), Cambridge and the Wellcome Sanger Institute This research was also supported by the NIHR Cambridge Biomedical Research Centre (BRC-1215-20014). The views expressed are those of the authors and not necessarily those of the NIHR or the Department of Health and Social Care.

## Figure legends

**Supplementary Figure 1.**
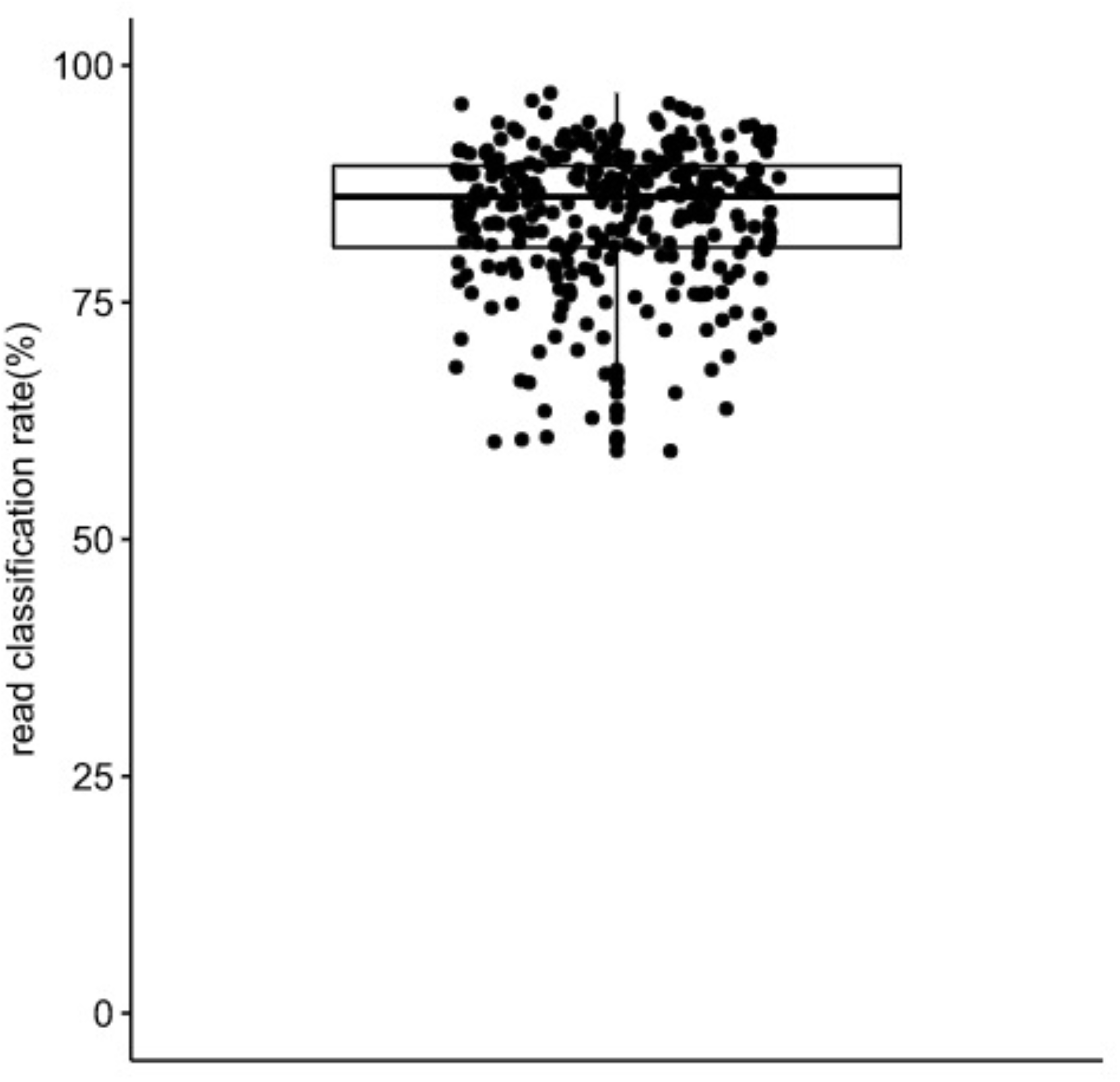
Shotgun sequencing read classification by our platform. Each sample was classified against a reference database and median read classification rate was 86%. Box and whiskers show, median and interquartile range.

**Supplementary Figure 2.**
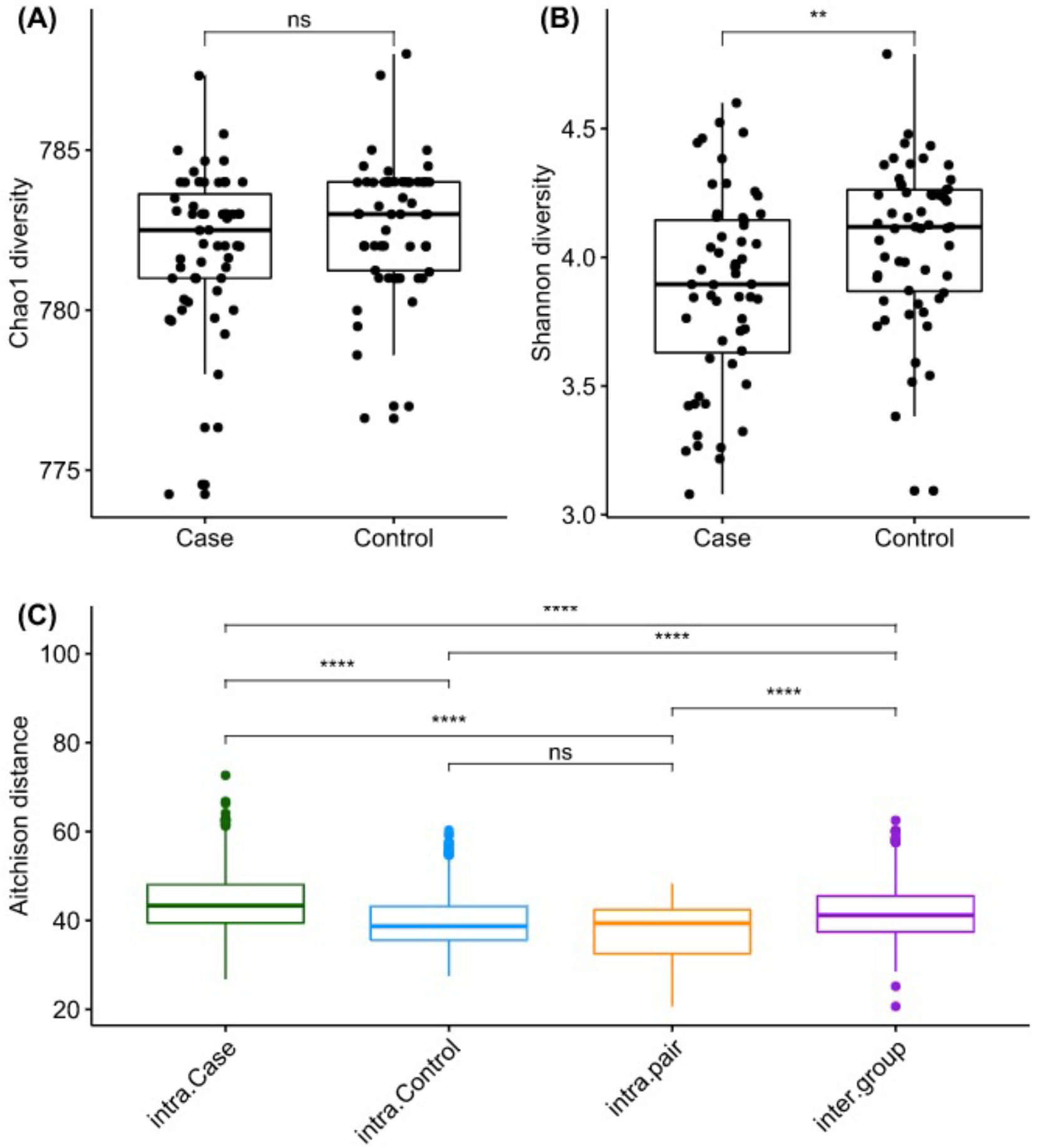
Alpha diversity analysis of microbiota profiles of cases and controls. Richness quantified by Chao1 diversity (A) and evenness measured by Shannon diversity (B) were computed for all case and control samples. Wilcoxon test was used to estimate statistical significance of the difference between groups. (C) Beta diversity analysis of microbiota profiles of cases and controls. Pairwise Aitchison distance was computed for all samples. Within-group variability was estimated for cases (green) and controls (blue). Intra-pair (orange) shows the variability measured for each household case-control pair. Inter-group distribution (purple) represents the comparison of any case and control samples in the absence of household pairs. Statistical difference between distributions are measured by Wilcoxon test p-values (****: p<0.0001, **: p<0.01, ns: p>0.05). Box and whiskers show, median and interquartile range.

**Supplementary Figure 3.**
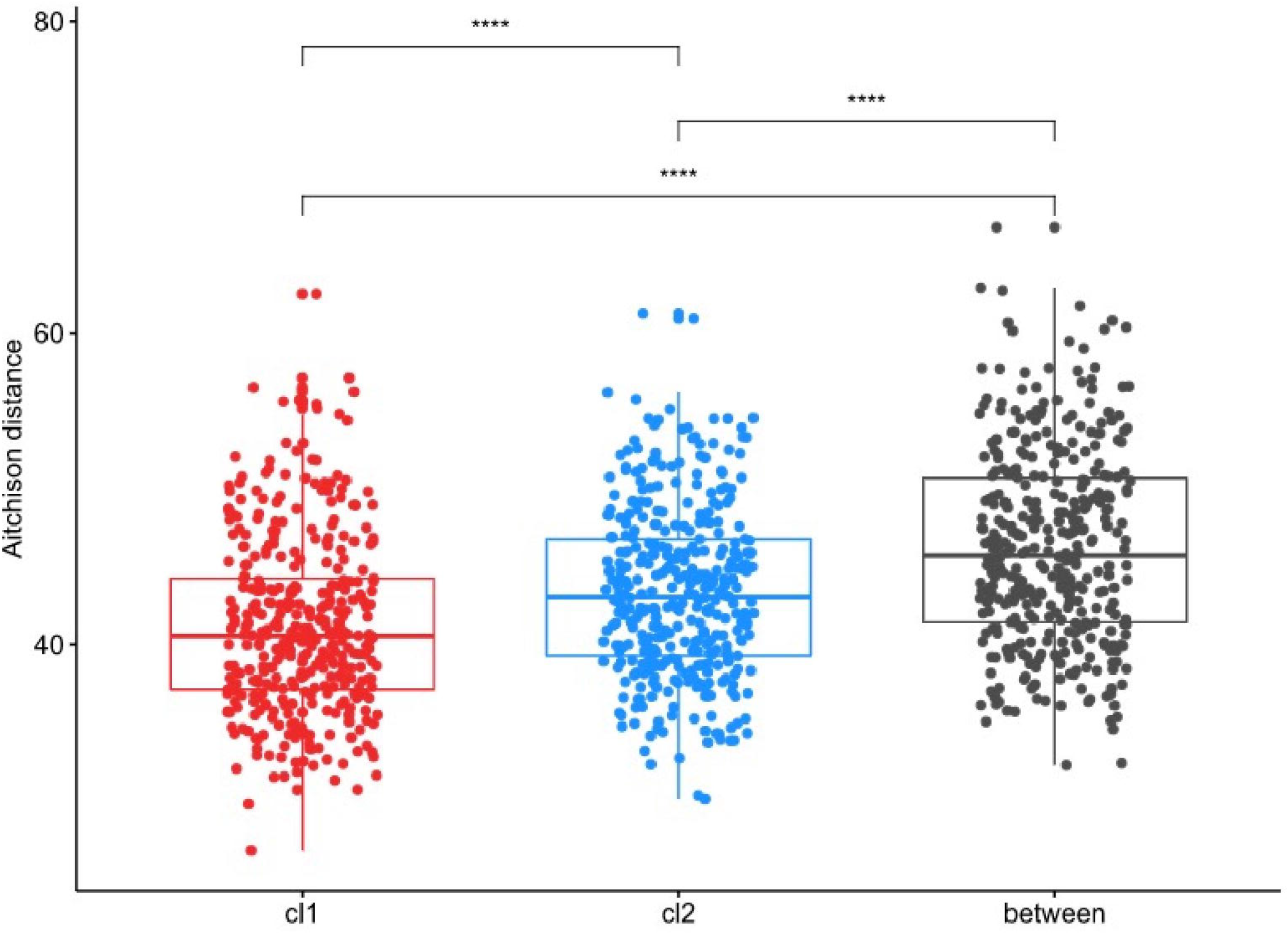
Beta diversity analysis of IBS cluster 1 (cl1) and cluster 2 (cl2) cases microbiota profiles. Pairwise Aitchison distance was computed for all samples. Within-group variability was estimated for IBS cluster 1 samples (red) and cluster 2 samples (blue). Between-group distribution (grey) represents the comparison of all IBS cluster 1 cases versus cluster 2 cases. Statistical difference between distributions are measured by Wilcoxon test p-values (****: p<0.0001). Box and whiskers show, median and interquartile range.

**Supplementary Figure 4.**
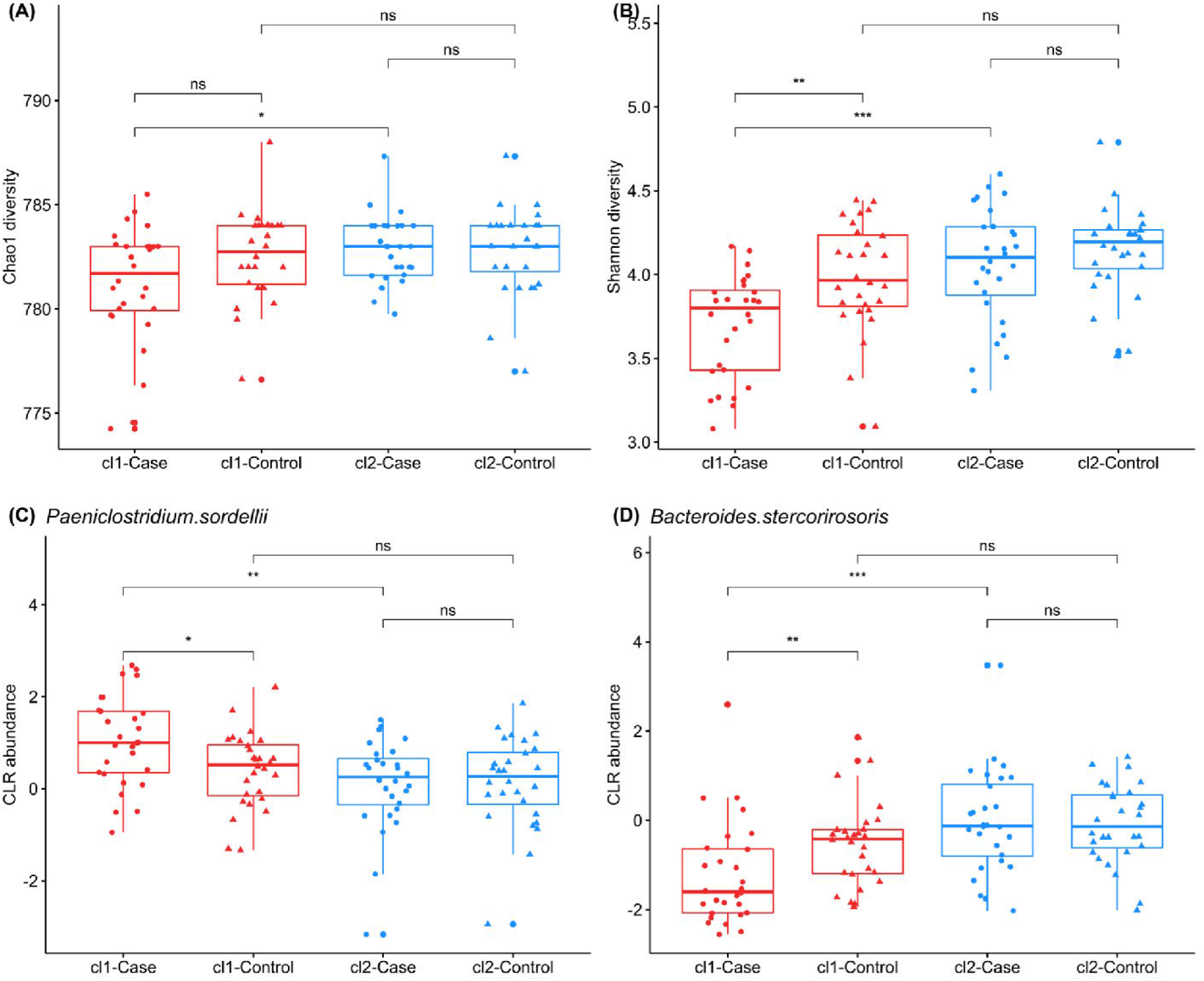
(A and B) Alpha diversity analysis of case (circles) and control (triangles) microbiota profiles after stratification by cluster 1 (cl1) and cluster 2 (cl2) sub-populations: richness quantified by Chao1 diversity (A) and evenness measured by Shannon diversity (B). (C and D) Compositional abundance analysis revealed key differences between 2 IBS subjects sub-populations. Center logratio (CLR) transformed abundance for representative species are shown. (C) Pathobiont species, such as Paeniclostridium sordelli, are more abundant in cluster 1 than in cluster 2 subjects. (D) Commensal species, such as Bacteroides genus members, are depleted in cluster 1 subjects when compared to cluster 2 subjects. Paired Wilcoxon test was used to estimate statistical significance of the difference between groups (***: p< 0.001, **: p<0.01, *:p<0.05, ns: p>0.05). Box and whiskers show, median and interquartile range.

**Supplementary Figure 5.**
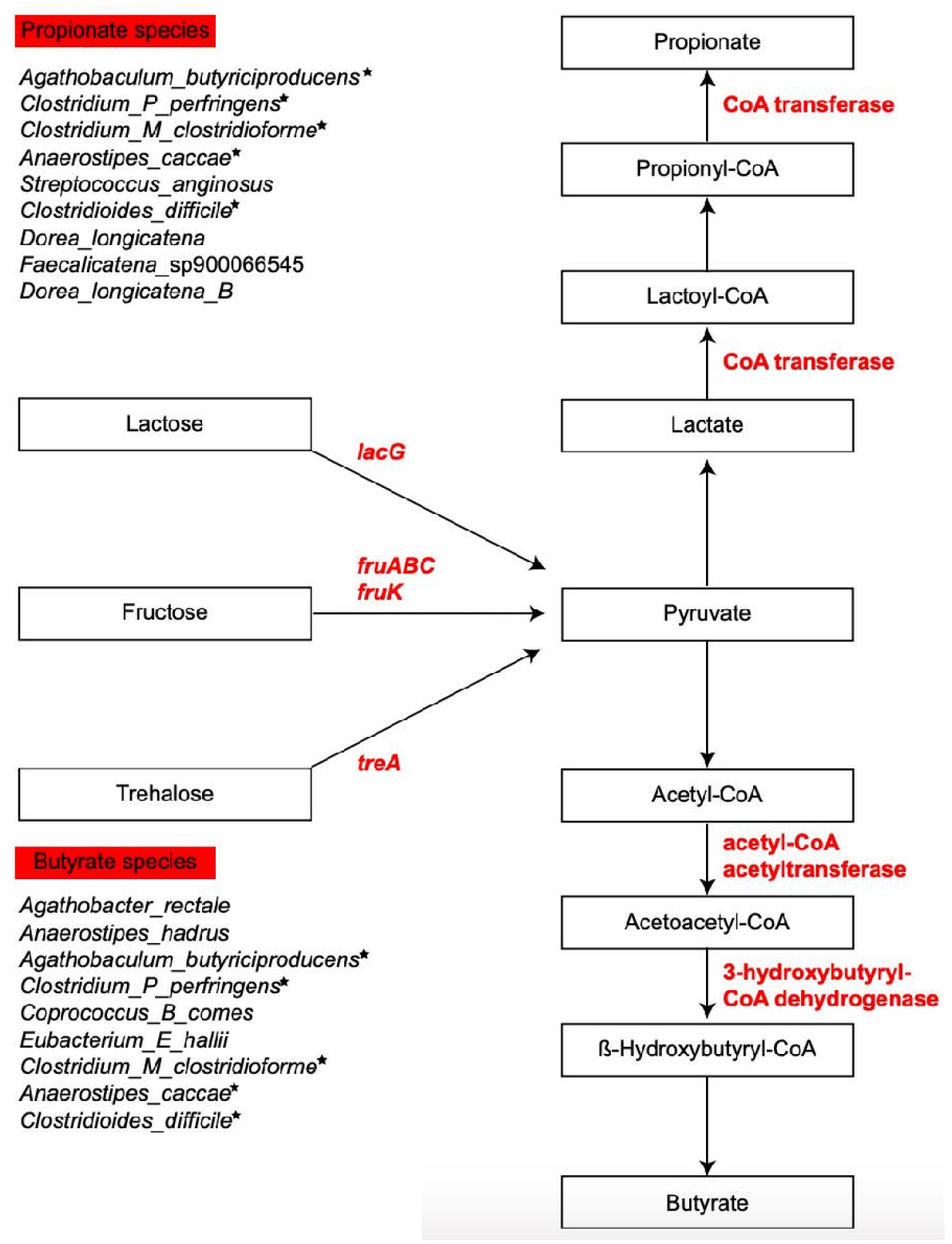
Pathway of simple dietary sugar metabolism and short chain fatty acid (butyrate and propionate) metabolism. Genes enriched in species representative of IBS^P^ are shown in red. Species that encode the enriched genes for butyrate and propionate metabolism are listed. Asterisks indicate species common between butyrate and propionate metabolism. lacG: 6-phospho-beta-galactosidase; fruABC: Putative PTS system system transporter subunit IIABC; fruK: 1-phosphofructokinase; treA: trehalose-6-phosphate hydrolase.

**Supplementary Figure 6.**
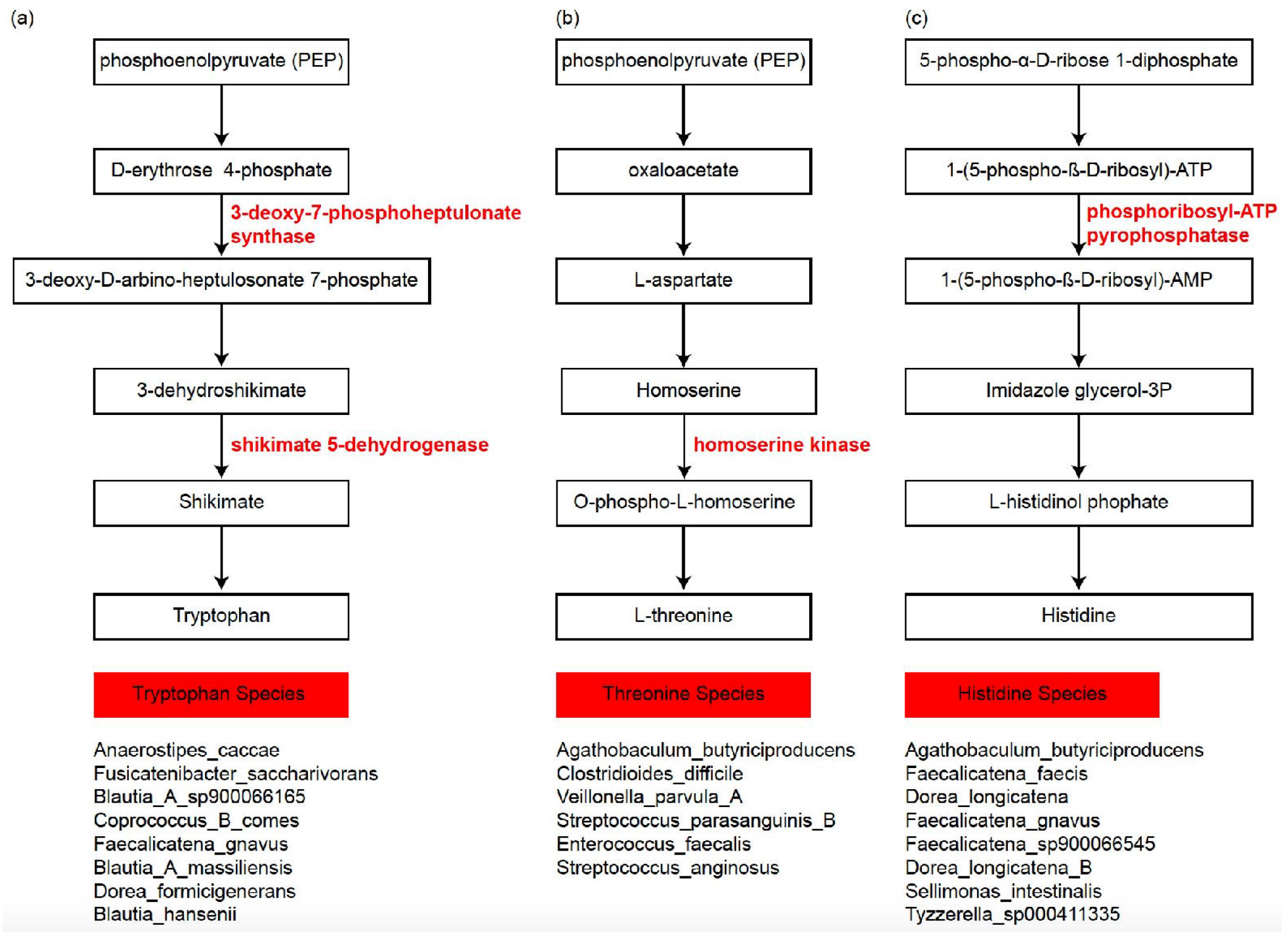
Pathway of tryptophan (a), threonine (b) and histidine (c) biosynthesis. Genes enriched in negatively associated species which are representative of IBS^P^ cluster are shown in red. Species that encode the enriched genes for tryptophan, threonine and histidine biosynthesis are listed.

**Supplementary Figure 7.**
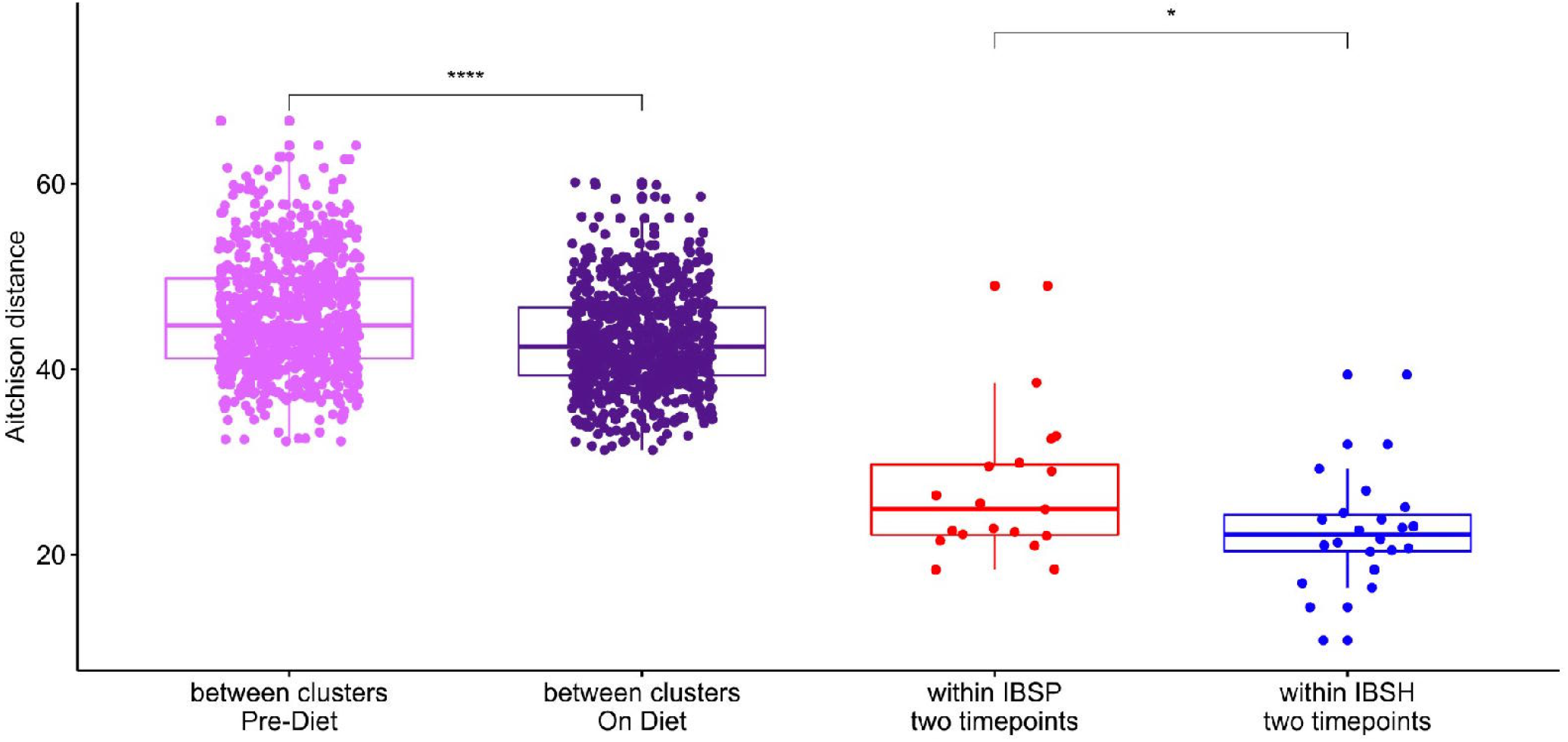
Pairwise Aitchison distance was computed for all case samples at pre-diet (light purple) and 4 weeks later on diet (dark purple). Longitudinal variability was estimated for IBS^P^ (red) and IBS^H^ (blue) samples, pairing samples from the same participant. Statistical difference between distributions are measured by Wilcoxon test p-values (***: p< 0.001, *:p<0.05). Box and whiskers show, median and interquartile range.

**Supplementary Figure 8.**
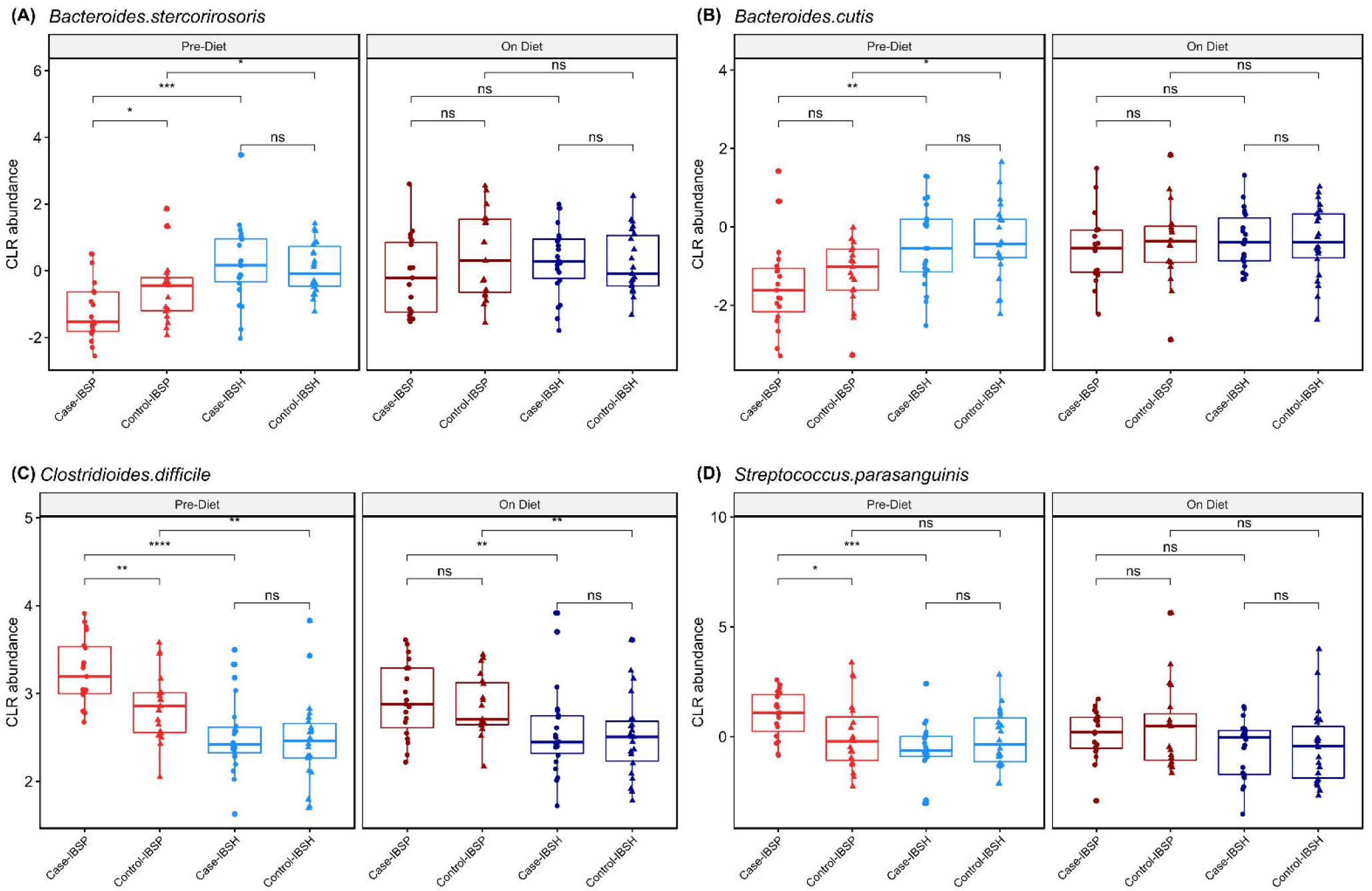
Diet intervention impact on taxonomic abundance for both case (circle) and matched control (triangle) samples. Linear mixed models identified differentially abundant species between IBS^P^ and IBS ^H^samples pre-diet and on diet intervention. (A, B) members of Bacteroides genus become more abundant in IBS^P^ during diet intervention (C,D) Pathobionts species become less abundant in IBS^P^ during diet intervention. Center logratio (CLR) transformed abundance for representative species are shown. Wilcoxon test was used to estimate statistical significance of the difference between groups (****: p<0.0001, ***: p<0.001, **: p<0.01, *:p<0.05, ns: p>0.05). Box and whiskers show, median and interquartile range.

**Supplementary Figure 9.**
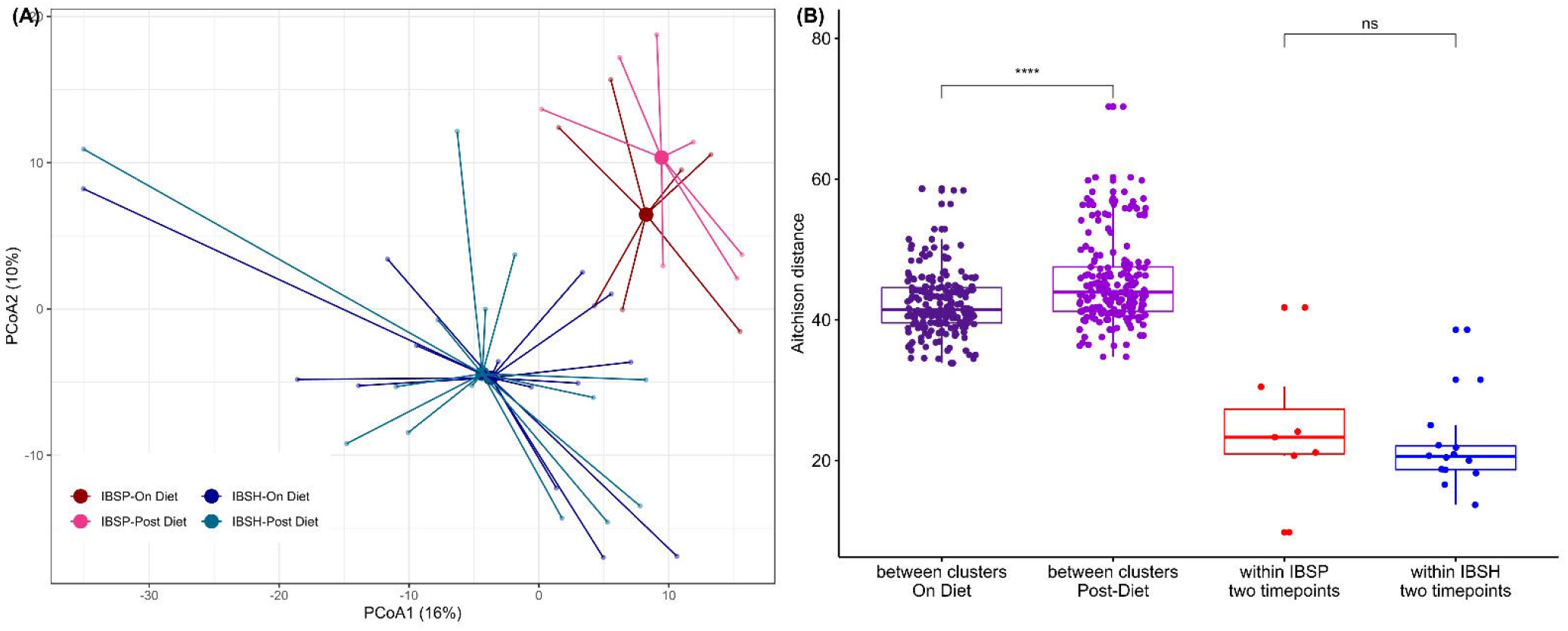
(A) Microbiota composition remains stable after cessation of the low FODMAP diet. Principal coordinate analysis of IBS^P^ and IBS^H^ subject sub-populations while on the low FODMAP dietary intervention and three months afterwards (by when cases either on normal diet or just excluding ‘trigger’ foods) projected on the two first components. (B) Pairwise Aitchison distance was computed for subjects at 4 weeks on-diet (dark purple) and 12 weeks post diet (purple). Longitudinal variability was estimated for IBS^P^ (red) and IBS^H^ (blue) samples, pairing samples from the same participant. Statistical difference between distributions are measured by Wilcoxon test p-values (****: p<0.0001, n.s.: p>0.05). Box and whiskers show, median and interquartile range.

## REFERENCES

1. Lovell RM, Ford AC. Global Prevalence of and Risk Factors for Irritable Bowel Syndrome: A Meta-analysis. Clinical Gastroenterology and Hepatology. 2012;10(7):712–21.

2. Simrén M, Svedlund J, Posserud I, Björnsson ES, Abrahamsson H. Health-related quality of life in patients attending a gastroenterology outpatient clinic: Functional disorders versus organic diseases. Clinical Gastroenterology and Hepatology. 2006;4(2):187–95.

3. Canavan C, West J, Card T. Review article: The economic impact of the irritable bowel syndrome. Alimentary Pharmacology and Therapeutics. 2014;40(9):1023–34.

4. Kuiken SD, Lindeboom R, Tytgat GN, Boeckxstaens GE. Relationship between symptoms and hypersensitivity to rectal distension in patients with irritable bowel syndrome. Alimentary Pharmacology and Therapeutics. 2005;22(2):157–64.

5. Camilleri M, Madsen K, Spiller R, Van Meerveld BG, Verne GN. Intestinal barrier function in health and gastrointestinal disease. Neurogastroenterology and Motility. 2012;24(6):503–12.

6. Vanuytsel T, Wanrooy SV, Vanheel H, Vanormelingen C, Houben E, Rasoel SS, et al. Psychological stress and corticotropin-releasing hormone increase intestinal permeability in humans by a mast cell-dependent mechanism. 2014.1293–9.

7. Thabane M, Kottachchi DT, Marshall JK. Systematic review and meta-analysis: The incidence and prognosis of post-infectious irritable bowel syndrome. Alimentary Pharmacology and Therapeutics. 2007;26(4):535–44.

8. Ford AC, Harris LA, Lacy BE, Quigley EMM, Moayyedi P. Systematic review with meta-analysis: the efficacy of prebiotics, probiotics, synbiotics and antibiotics in irritable bowel syndrome. Alimentary Pharmacology and Therapeutics. 2018;48(10):1044–60.

9. Dear KLE, Elia M, Hunter JO. Do interventions which reduce colonic bacterial fermentation improve symptoms of irritable bowel syndrome? Digestive Diseases and Sciences. 2005;50(4):758–66.

10. El-Salhy M, Hatlebakk JG, Gilja OH, Bråthen Kristoffersen A, Hausken T. Efficacy of faecal microbiota transplantation for patients with irritable bowel syndrome in a randomised, double-blind, placebo-controlled study. Gut. 2020;69(5):859–67.

11. Duan R, Zhu S, Wang B, Duan L. Alterations of Gut Microbiota in Patients With Irritable Bowel Syndrome Based on 16S rRNA-Targeted Sequencing: A Systematic Review. Clin Transl Gastroenterol. 2019;10(2):e00012.

12. Jeffery IB, O’Toole PW, Öhman L, Claesson MJ, Deane J, Quigley EMM, et al. An irritable bowel syndrome subtype defined by species-specific alterations in faecal microbiota. Gut. 2012;61(7):997–1006.

13. Jeffery IB, Das A, O’Herlihy E, Coughlan S, Cisek K, Moore M, et al. Differences in Fecal Microbiomes and Metabolomes of People With vs Without Irritable Bowel Syndrome and Bile Acid Malabsorption. Gastroenterology. 2020;158(4):1016–28 e8.

14. Chumpitazi BP, Cope JL, Hollister EB, Tsai CM, McMeans AR, Luna RA, et al. Randomised clinical trial: gut microbiome biomarkers are associated with clinical response to a low FODMAP diet in children with the irritable bowel syndrome. Alimentary Pharmacology and Therapeutics. 2015;42(4):418–27.

15. Halmos EP, Power VA, Shepherd SJ, Gibson PR, Muir JG. A Diet Low in FODMAPs Reduces Symptoms of Irritable Bowel Syndrome. Gastroenterology. 2014;146(1):67-75.e5.

16. Staudacher HM, Lomer MCE, Farquharson FM, Louis P, Fava F, Franciosi E, et al. A Diet Low in FODMAPs Reduces Symptoms in Patients With Irritable Bowel Syndrome and A Probiotic Restores Bifidobacterium Species: A Randomized Controlled Trial. Gastroenterology. 2017;153(4):936–47.

17. Whelan K, Martin LD, Staudacher HM, Lomer MCE. The low FODMAP diet in the management of irritable bowel syndrome: an evidence-based review of FODMAP restriction, reintroduction and personalisation in clinical practice. Journal of Human Nutrition and Dietetics. 2018;31(2):239–55.

18. Chumpitazi BP. The gut microbiome as a predictor of low fermentable oligosaccharides disaccharides monosaccharides and polyols diet efficacy in functional bowel disorders. Current Opinion in Gastroenterology. 2020;36(2):147–54.

19. Gibson PR, Halmos EP, Muir JG. Review article: FODMAPS, prebiotics and gut health-the FODMAP hypothesis revisited. Alimentary Pharmacology and Therapeutics. 2020;52(2):233–46.

20. Chang L, Di Lorenzo C, Farrugia G, Hamilton FA, Mawe GM, Pasricha PJ, et al. Functional Bowel Disorders: A Roadmap to Guide the Next Generation of Research. Gastroenterology. 2018;154(3):723–35.

21. Byrd AL, Liu M, Fujimura KE, Lyalina S, Nagarkar DR, Charbit B, et al. Gut microbiome stability and dynamics in healthy donors and patients with non-gastrointestinal cancers. J Exp Med. 2021;218(1).

22. Morais LH, Schreiber HLt, Mazmanian SK. The gut microbiota-brain axis in behaviour and brain disorders. Nat Rev Microbiol. 2021;19(4):241–55.

23. Zheng D, Liwinski T, Elinav E. Interaction between microbiota and immunity in health and disease. Cell Res. 2020;30(6):492–506.

24. Forster SC, Kumar N, Anonye BO, Almeida A, Viciani E, Stares MD, et al. A human gut bacterial genome and culture collection for improved metagenomic analyses. Nat Biotechnol. 2019;37(2):186–92.

25. Palsson OS, Whitehead WE, Van Tilburg MAL, Chang L, Chey W, Crowell MD, et al. Development and validation of the Rome IV diagnostic questionnaire for adults. Gastroenterology. 2016;150(6):1481–91.

26. Imhann F, Vich Vila A, Bonder MJ, Lopez Manosalva AG, Koonen DPY, Fu J, et al. The influence of proton pump inhibitors and other commonly used medication on the gut microbiota. Gut microbes. 2017;8(4):351–8.

27. Francis CY, Morris J, Whorwell PJ. The irritable bowel severity scoring system: a simple method of monitoring irritable bowel syndrome and its progress. Alimentary Pharmacology & Therapeutics. 1997;11(2):395–402.

28. Lu J, Breitwieser FP, Thielen P, Salzberg SL. Bracken: Estimating species abundance in metagenomics data. PeerJ Computer Science. 2017;2017(1):1–17.

29. R Core Team. R: A language and environment for statistical computing.. 2020.

30. Gloor GB, Macklaim JM, Pawlowsky-Glahn V, Egozcue JJ. Microbiome datasets are compositional: And this is not optional. Frontiers in Microbiology. 2017;8(NOV):1–6.

31. Palarea-Albaladejo J, Martin-Fernandez J. zCompositions – R package for multivariate imputation of left-censored data under a compositional approach. Chemom Intell Lab Syst. 2015;143:85--96.

32. Oksanen J, Kindt R, Legendre P, O’Hara B, Simpson GL, Solymos P, et al. vegan: Community Ecology Package. R package version 2.5-7 ed2020.

33. Anderson MJ. A new method for non-parametric multivariate analysis of variance. Austral Ecology. 2001;26(1):32–46.

34. Mallick H, Rahnavard A, McIver LJ, Ma S, Zhang Y, Nguyen LH, et al. Multivariable Association Discovery in Population-scale Meta-omics Studies. bioRxiv. 2021:2021.01.20.427420.

35. Zou Y, Xue W, Luo G, Deng Z, Qin P, Guo R, et al. 1,520 reference genomes from cultivated human gut bacteria enable functional microbiome analyses. Nat Biotechnol. 2019;37(2):179–85.

36. Price MN, Dehal PS, Arkin AP. FastTree 2--approximately maximum-likelihood trees for large alignments. PLoS One. 2010;5(3):e9490.

37. Chaumeil PA, Mussig AJ, Hugenholtz P, Parks DH. GTDB-Tk: a toolkit to classify genomes with the Genome Taxonomy Database. Bioinformatics. 2019.

38. Letunic I, Bork P. Interactive Tree Of Life (iTOL) v4: recent updates and new developments. Nucleic Acids Res. 2019;47(W1):W256–W9.

39. Franzosa EA, McIver LJ, Rahnavard G, Thompson LR, Schirmer M, Weingart G, et al. Species-level functional profiling of metagenomes and metatranscriptomes. Nat Methods. 2018;15(11):962–8.

40. Caspi R, Billington R, Keseler IM, Kothari A, Krummenacker M, Midford PE, et al. The MetaCyc database of metabolic pathways and enzymes - a 2019 update. Nucleic Acids Res. 2020;48(D1):D445–D53.

41. UniProt C. UniProt: the universal protein knowledgebase in 2021. Nucleic Acids Res. 2021;49(D1):D480–D9.

42. Altschul SF, Gish W, Miller W, Myers EW, Lipman DJ. Basic local alignment search tool. J Mol Biol. 1990;215(3):403–10.

43. Browne HP, Neville BA, Forster SC, Lawley TD. Europe PMC Funders Group Transmission of the gut microbiota : spreading of health. Nature Reviews Microbiology. 2018;15(9):531–43.

44. Hall AB, Yassour M, Sauk J, Garner A, Jiang X, Arthur T, et al. A novel Ruminococcus gnavus clade enriched in inflammatory bowel disease patients. Genome Med. 2017;9(1):103.

45. Vujkovic-Cvijin I, Sklar J, Jiang L, Natarajan L, Knight R, Belkaid Y. Host variables confound gut microbiota studies of human disease. Nature. 2020;587(7834):448–54.

46. Collins J, Robinson C, Danhof H, Knetsch CW, van Leeuwen HC, Lawley TD, et al. Dietary trehalose enhances virulence of epidemic Clostridium difficile. Nature. 2018;553(7688):291–4.

47. Macfarlane GT, Macfarlane S. Bacteria, colonic fermentation, and gastrointestinal health. J AOAC Int. 2012;95(1):50–60.

48. Koh A, De Vadder F, Kovatcheva-Datchary P, Backhed F. From Dietary Fiber to Host Physiology: Short-Chain Fatty Acids as Key Bacterial Metabolites. Cell. 2016;165(6):1332–45.

49. Reigstad CS, Salmonson CE, Rainey JF, 3rd, Szurszewski JH, Linden DR, Sonnenburg JL, et al. Gut microbes promote colonic serotonin production through an effect of short-chain fatty acids on enterochromaffin cells. FASEB J. 2015;29(4):1395–403.

50. Yano JM, Yu K, Donaldson GP, Shastri GG, Ann P, Ma L, et al. Indigenous bacteria from the gut microbiota regulate host serotonin biosynthesis. Cell. 2015;161(2):264–76.

51. Lee JS, Kim SY, Chun YS, Chun YJ, Shin SY, Choi CH, et al. Characteristics of fecal metabolic profiles in patients with irritable bowel syndrome with predominant diarrhea investigated using (1) H-NMR coupled with multivariate statistical analysis. Neurogastroenterol Motil. 2020;32(6):e13830.

52. Bennet SMP, Böhn L, Störsrud S, Liljebo T, Collin L, Lindfors P, et al. Multivariate modelling of faecal bacterial profiles of patients with IBS predicts responsiveness to a diet low in FODMAPs. Gut. 2018;67(5):872–81.

53. Rossi M, Aggio R, Staudacher HM, Lomer MC, Lindsay JO, Irving P, et al. Volatile Organic Compounds in Feces Associate With Response to Dietary Intervention in Patients With Irritable Bowel Syndrome. Clinical Gastroenterology and Hepatology. 2018;16(3):385-91.e1.

